# PACSIN2-dependent apical endocytosis regulates the morphology of epithelial microvilli

**DOI:** 10.1101/686816

**Authors:** Meagan M. Postema, Nathan E. Grega-Larson, Leslie M. Meenderink, Matthew J. Tyska

## Abstract

Apical microvilli are critical for the homeostasis of transporting epithelia, yet mechanisms that control the assembly and morphology of these protrusions remain poorly understood. Previous studies in intestinal epithelial cell lines suggested a role for F-BAR domain protein PACSIN2 in normal microvillar assembly. Here we report the phenotype of PACSIN2 KO mice and provide evidence that through its role in promoting apical endocytosis, this molecule functions in controlling microvillar morphology. PACSIN2 KO enterocytes exhibit reduced numbers of microvilli and defects in microvillar ultrastructure, with membranes lifting away from rootlets of core bundles. Dynamin2, a PACSIN2 binding partner, and other endocytic factors were also lost from their normal localization near microvillar rootlets. To determine if loss of endocytic machinery could explain defects in microvillar morphology, we examined the impact of PACSIN2 KD and endocytosis inhibition on live intestinal epithelial cells. These assays revealed that when endocytic vesicle scission fails, tubules are pulled into the cytoplasm and this, in turn, leads to a membrane lifting phenomenon reminiscent of that observed in PACSIN2 KO brush borders. These findings lead to a new model where inward forces generated by endocytic machinery on the plasma membrane control the membrane wrapping of cell surface protrusions.

**Highlight for TOC:** Apical microvilli increase the functional surface area of transporting epithelia. Here we report that the F-BAR domain-containing protein PACSIN2, through its ability to promote apical endocytosis, plays a critical role in controlling the morphology of intestinal brush border microvilli.

## INTRODUCTION

Apical specializations enable epithelial cells to carry out specific functions including solute uptake and mechano-sensation. In the context of transporting epithelia, the apical surface is occupied by actin bundle-supported microvilli: finger-like protrusions that serve to amplify membrane surface area and maximize solute uptake capacity [1]. A well-studied example is found in the intestinal tract where enterocytes, the most abundant epithelial cell type in the gut, provide the sole site of nutrient absorption. Enterocytes build tightly-packed arrays of 1000s of microvilli, known as a brush borders. Microvillar growth and ordered packing take place as enterocytes differentiate, which occurs as they exit stem cell-containing crypt domains and move onto the villus surface [2–4].

Microvillus formation requires coordination of a variety of activities, including actin filament nucleation, elongation, and bundling, which presumably all occur at the interface with the apical plasma membrane. Nucleation of the actin filaments that comprise core bundles is at least partially controlled by the WH2 domain protein cordon bleu (COBL), which is required for normal brush border assembly in intestinal epithelial cell lines [5–7]. COBL overexpression drives microvillus elongation and also leads to protrusions that are straighter, with higher actin content [6]. COBL localizes to microvillar rootlets, which are embedded in a dense sub-apical network of intermediate filaments known as the terminal web [8]. The actin bundling protein fimbrin also localizes to the terminal web and has been shown to link microvillar actin to keratin-19 in intermediate filament [9]. Along with fimbrin, two other bundling proteins, villin and espin, stabilize the core bundle in a region-specific manner [10–13] and may promote elongation by slowing disassembly at the pointed ends [14]. Later in differentiation, epithelial-specific protocadherins target to the tips of microvilli to promote their elongation and tight packing [15–20]. Such intermicrovillar adhesion allows cells to generate the maximum number of protrusions per unit apical surface area [21].

Another recently identified factor that functions in microvillar growth is the I-BAR (inverse-Bin-Amphiphysin-Rvs) domain containing protein, insulin receptor tyrosine kinase substrate (IRTKS) [22]. BAR domains are small, three helix bundles that form curved dimers ~20 nm in length, which in turn form higher order oligomers capable of sensing and inducing membrane curvature [23]. I-BAR domains exhibit a structural curvature that is well matched to membrane bending away from the cell [24], like that found at the distal tips of microvilli. Indeed, IRTKS targets to microvillar tips where it promotes elongation directly by interacting with the core actin bundle, and indirectly through its interactions with epidermal growth factor receptor pathway substrate 8 (EPS8), another tip targeting factor implicated in the elongation of finger-like protrusions [22, 25-28].

In contrast to the curvature preference of I-BAR domains, F-BAR (Fes-CIP4 homology Bin-amphiphysin-Rvs161/167) motifs prefer binding to membranes that curve in toward the cytoplasm [29–31]. Protein kinase C and casein kinase substrate in neurons (PACSIN) family members are F-BAR proteins that have been implicated in a variety of cellular processes, including clathrin-dependent and independent endocytosis, caveolae formation, vesicle trafficking, actin dynamics, and cell migration [32–38]. Although PACSIN2 is widely expressed [39], PACSIN1 exhibits specificity for neural tissues [40], whereas PACSIN3 is expressed in heart and skeletal muscle [41]. All three PACSIN isoforms contain an N-terminal F-BAR domain, along with a C-terminal SH3 domain. Interestingly, previous studies in intestinal epithelial cells revealed that PACSIN2 localizes to the intermicrovillar region in the terminal web, which exhibits a high degree of inward bending and also serves as the site of endocytosis [6]. Moreover, through its SH3 domain, PACSIN2 also interacts with several binding partners with roles in actin filament nucleation and endocytosis at the membrane-cytoskeleton interface (Fig 1A). One example is the actin nucleator COBL, which interacts with PACSIN2 in the terminal web. Loss-of-function studies in intestinal epithelial cell lines suggest that PACSIN2 serves to recruit or anchor COBL in this location [6]. COBL in turn uses its multiple WH2 domains to promote elongation of core actin bundles [7]. In this context, PACSIN2 is critical for normal microvillar growth as knocking down the molecule in cell culture models leads to defects in brush border assembly [6]. A second SH3 binding partner that links PACSIN2 to actin assembly is N-WASP, a nucleation promoting factor and adaptor protein that activates the ubiquitous branched actin nucleator, ARP2/3 [42]. N-WASP interactions with PACSIN2 are believed to physically link the actin cytoskeleton to membranes in processes such as endocytosis [43, 44]. Yet another link between PACSIN2 and endocytosis is mediated by SH3 domain binding to the large GTPase Dynamin2, which drives vesicle excision from the plasma membrane. PACSIN2 binds to and recruits Dynamin2 in the context of clathrin-mediated endocytosis and the internalization of caveolae [45]. Other studies have shown that the F-BAR domain of PACSIN2 is capable of oligomerizing and coating the necks of newly forming vesicles, which likely helps stabilize these intermediates before excision [46].

**Figure 1.**
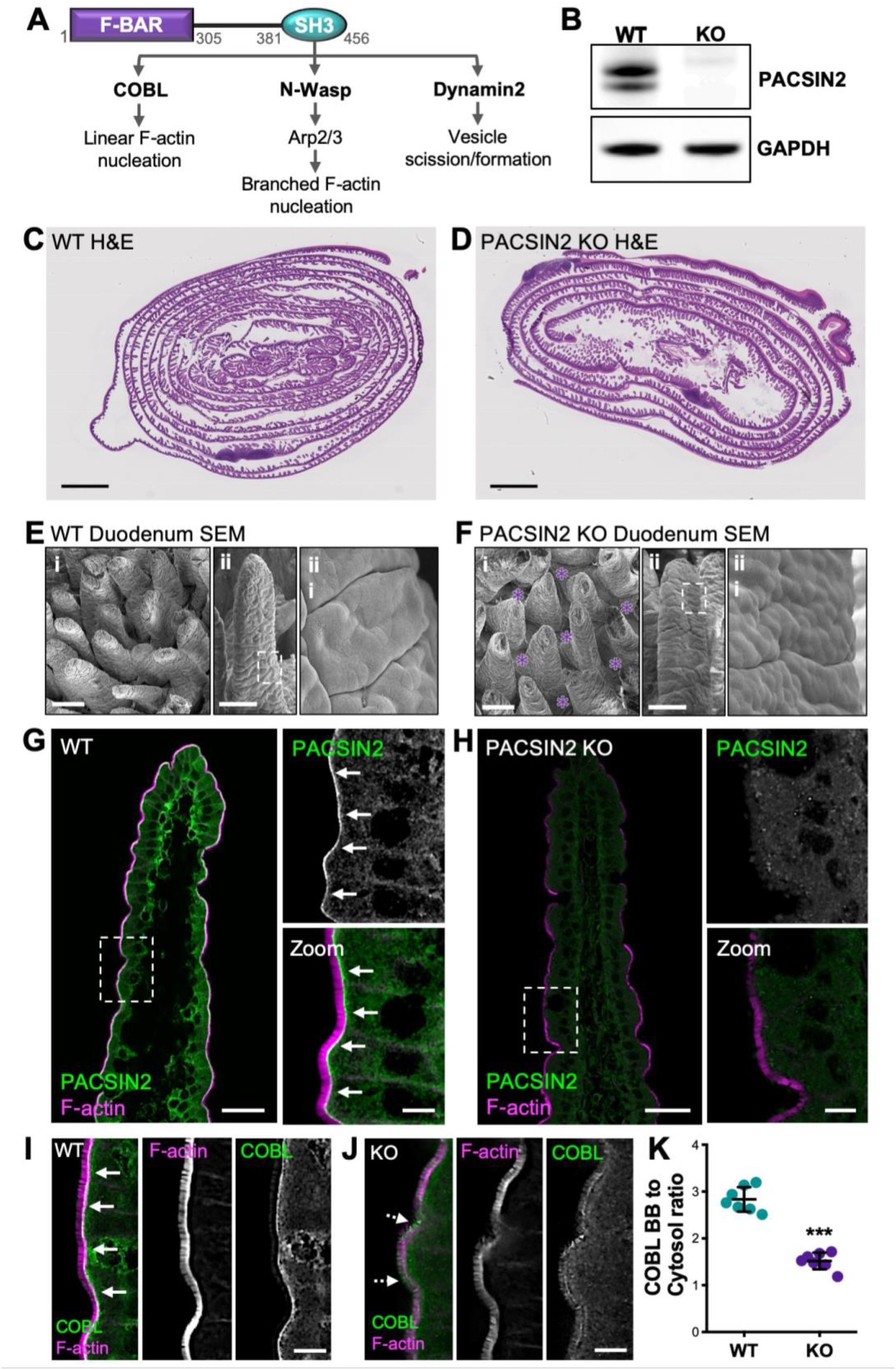
PACSIN2 KO disrupts COBL localization. **(A)** PACSIN2 domain diagram depicting SH3 binding partners and prospective functions. **(B)**Western blot of WT and PACSIN2 KO tissue with GAPDH as a loading control. **(C, D)** H&E-stained swiss roll sections of paraffin-embedded small intestine from WT and PACSIN2 KO mice. Scale bars, 2mm. **(E, F)** Scanning EM images of intestinal tissue samples from WT (E) and PACSIN2 KO (F) mice. Scale bars, 100 μm for *i*, 100 μm for *ii*, and 10 μm for *iii*; purple asterisks in KO *Bi* indicate bare spaces in the epithelium between adjacent villi. **(G, H)** Endogenous PACSIN2 (green) and phalloidin (F-actin, magenta) labelling of WT and PACSIN2 KO frozen tissue sections. Arrows highlight PACSIN2 signal at the base of the brush border in WT tissue (G). Scale bars, 50 μm for main panels, 10 μm for zooms. **(I, J)** Endogenous COBL (green) and phalloidin (magenta) labelling of WT and PACSIN2 KO frozen tissue sections. Solid arrows highlight COBL signal at the base of the brush border in WT tissue (I), dashed arrows highlight mislocalization of COBL signal in KO tissue (J). Scale bars, 10 μm. **(K)** Quantification of the ratio of COBL brush border (BB) to cytosol signal intensity between the WT and PACSIN2 KO tissue; n = 7 tissue sections per condition. Error bars indicate ± SD; p value was calculated using a t test (***p<0.001).

In the present study, we sought to develop our understanding of PACSIN2 function in the epithelial apical domain through the analysis of mice lacking PACSIN2 expression. Ultrastructural studies of tissues from KO animals revealed a plasma membrane lifting phenotype, where core actin bundles are no longer fully enveloped in membrane, and in some cases fuse with adjacent protrusions. Moreover, Dynamin2 and other endocytic factors were lost from their normal localization near the intermicrovillar endocytic region. To determine if the loss of endocytic machinery could explain defects in brush border morphology, we examined the impact of dynamin inhibition and PACSIN2 KD on live intestinal epithelial cells. We found that when endocytic vesicle scission fails, tubules are pulled into the cytoplasm, and this leads directly to a membrane lifting phenomenon similar to that observed in PACSIN2 KO brush borders. Our findings illuminate a previously unrecognized link between endocytic function and the morphology of the epithelial apical domain, and also suggest that inward forces generated on the plasma membrane by endocytic machinery control the membrane wrapping of cell surface protrusions.

## RESULTS

### PACSIN2 KO disrupts COBL localization

To explore how PACSIN2 contributes to enterocyte apical architecture and brush border assembly *in vivo*, we acquired mice expressing a PACSIN2^tm1b(EUCOMM)Hmgu^ allele from the KOMP resource [47]. Tm1b mice are “CREed knockout first” and provide constitutive loss of expression in all tissues. KO of PACSIN2 was confirmed using western blot analysis (Fig.1B). PACSIN2 KO mice did not exhibit gross level phenotypes or defects in growth. Analysis of hematoxylin- and eosin-stained swiss roll sections (Fig. 1C,D) and scanning electron microscopy (SEM) of duodenal tissue sections (Fig. 1E,F) revealed that PACSIN2 KO tissues were morphologically similar to WT. In frozen sections of WT tissue, PACSIN2 is strongly enriched at the base of the brush border in the terminal web (Fig. 1G). However, this labeling is completely lost in KO mice, further confirming loss of expression (Fig. 1H). As previous studies in intestinal epithelial cells lines suggest that this F-BAR protein functions in the recruitment of COBL, we next sought to determine if COBL was mislocalized in the absence of PACSIN2. As expected, COBL exhibits high level enrichment in the terminal web of WT tissues (Fig. 1I), but this labeling is significantly perturbed in KO samples (Fig. 1J). This point was also confirmed with quantification of brush border to cytosol intensity ratios, which were markedly reduced in KO samples (2.83 ± 0.26 WT vs. 1.52 ± 0.18 KO; Fig. 1K). Interestingly, in KO tissues COBL signal also appears redistributed along the microvillar axis (dashed arrows, Fig. 1J), suggesting a role for PACSIN2 in anchoring COBL near microvillar rootlets. These results confirm the loss of PACSIN2 in the KO intestinal tissue and are consistent with previous studies indicating that PACSIN2 is needed for efficient targeting of COBL to the apical domain.

### Loss of PACSIN2 decreases apical and basolateral F-actin levels

Given that PACSIN2 and its binding partners have been implicated in actin network assembly [35, 48], we next sought to determine if KO tissues exhibited perturbations in the actin cytoskeleton. Indeed, our initial staining of KO frozen tissue sections (Fig. 1H) suggested that apical F-actin levels (assessed with phalloidin staining) were reduced, especially in the distal regions of villi. To examine this in greater detail, we performed volumetric imaging of whole mounted segments of intestinal tissue stained for F-actin. 3D reconstructions of individual villi revealed several striking defects in KO samples (Fig. 2A,B). Levels of F-actin appeared reduced throughout the apical domain, both in the brush border and at the lateral margins of cells (Fig. 2A,B). We also noted that the apical surfaces of individual cells exhibited a domed appearance, curving outward towards the lumen (Fig. 2B). This phenotype was even more evident when we examined the apical surface of KO tissues using SEM (Fig. 1E,F). In higher-resolution tilted 3D projections, KO brush borders demonstrated an apparent thinning of the F-actin signal, with certain regions exhibiting significantly reduced microvillar density relative to WT controls (Fig. 2C,E). Line-scans drawn through the single plane images (Fig. 2C,E, bottom panels) showed an almost 2-fold decrease in the PACSIN2 KO F-actin signal with several gaps throughout (maximum F-actin peak signal of 4095 for WT vs. 2156 for KO; Fig. 2D,F). Further quantification using thresholding and segmentation on multiple tissue sections also indicated a marked decrease in brush border F-actin intensity in PACSIN2 KO tissues (mean intensity units 1912 ± 323 for WT vs. 1123 ± 239 for KO; Fig. 2G-I). Together these data indicate that in the absence of PACSIN2, actin polymerization at the apical surface is compromised.

**Figure 2.**
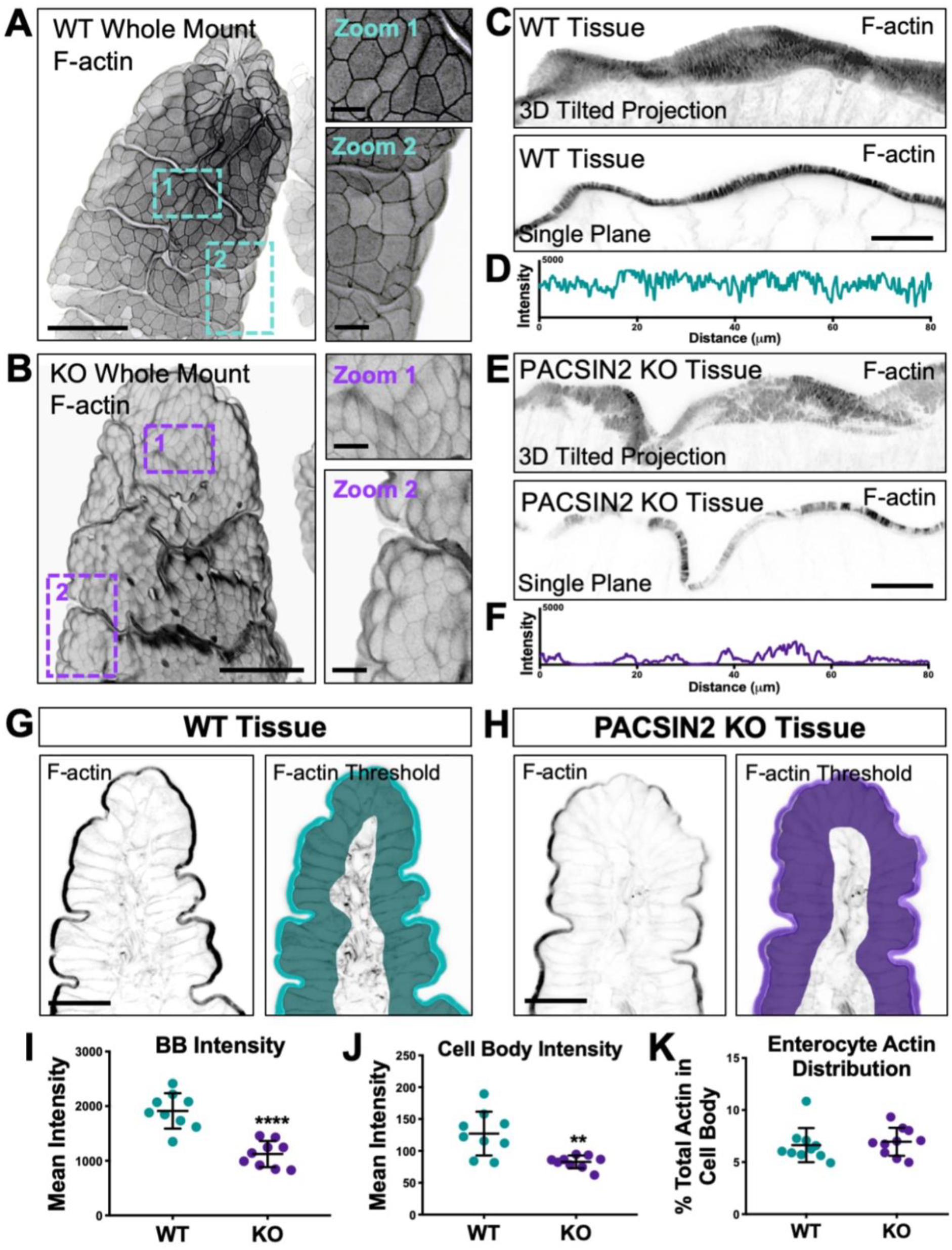
Loss of PACSIN2 decreases apical and basolateral F-actin levels. **(A, B)** 3D projections of 50 μm sections of WT (A) and PACSIN2 KO (B) whole mount tissue. Zooms highlight differences in cell surface morphology and actin intensity between WT and KO. Actin signal is inverted to simplify visualization; Scale bars, 50 μm for main panels, 10 μm for zooms. (C, E) 3D reconstructed volumes of 8 μm sections (top) and single image planes (bottom) of phalloidin stained WT and PASCIN2 KO frozen tissue sections. Scale bars, 25 μm. (D, F) Plots of raw 8-bit intensity data from an 80 μm line drawn through the brush border of the single plane images. PACSIN2 KO tissue has ~2-fold decrease in brush border actin intensity. (G, H) Phalloidin labelling of WT and PACSIN2 KO frozen tissue sections. Right panels show representative thresholding of brush border and cell body used in quantification (I-K). Scale bars, 50 μm. **(I)** Quantification of brush border (BB) actin intensity between WT and PACSIN2 KO tissue; 9 tissue sections per condition. **(J)** Quantification of cell body actin intensity between the WT and PACSIN2 KO tissue; 9 tissue sections per condition. **(K)** Quantification of the percent of actin in the cell body to total actin between WT and PACSIN2 KO; 9 tissue sections per condition. Error bars indicate ± SD; p values were calculated using a t test (**p<0.01, ****p<0.0001).

Given the striking reduction of apical F-actin signal observed in PACSIN2 KO brush borders, we also examined F-actin levels in actin networks in other parts of the cell (Fig. 2G,H). Mean F-actin intensity values, measured using a threshold that included all cellular structures basolateral to the brush border, were also markedly reduced (127.2 ± 34.5 WT vs. 82.7 ± 10.0 KO; Fig. 2J). Interestingly, ratios of brush border/cell body F-actin intensities were unchanged in KO relative to WT samples (Fig. 2K), suggesting that the overall distribution of actin polymer was similar. Further analysis of the cell body F-actin signal revealed that most of the intensity is derived from the basolateral margins, at sites of cell-cell contact (Fig. S1E-J). Linescan analysis through multiple cells revealed that junctional F-actin levels were also significantly reduced at these sites (Fig. S1G,J). Consistent with this, we also noted defects in the localization of tight and adherens junction markers, ZO-1 and E-cadherin; both probes exhibited significantly lower levels of junctional enrichment relative to WT tissue sections (Fig. S1K-M). These data indicate that in addition to promoting the growth of microvilli on the apical surface, PACSIN2 also drives the accumulation of F-actin at cell margins, where it promotes accumulation of factors that contribute to cell-cell adhesion.

### Endocytic machinery is mislocalized in the absence of PACSIN2

In addition to scaffolding factors such as COBL and N-WASP, which promote actin polymerization in the apical and basolateral compartments, PACSIN2 has also been implicated in endocytic function in epithelial cells. Therefore, we sought to determine if the sub-apical endocytic compartment in the terminal web was disrupted in the absence of PACSIN2. Under normal conditions, Dynamin2 is highly enriched at the base of microvilli in the terminal web, the site of endocytic vesicle formation and fission (Fig. 3A). However, upon KO of PACSIN2, this striking band of enrichment is lost (Fig. 3B), which is also reflected in a significant decrease of the brush border to cytosol ratio for this signal (2.04 ± 0.76 WT vs. 1.20 ± 0.25 KO; Fig. 3C). We also stained sections for VAMP4 (vesicle associated membrane protein 4), which has established roles in endo- and exocytosis [49, 50]. Similar to Dynamin2, VAMP4 exhibits striking enrichment at the base of the brush border in WT samples (Fig. 3D), but marked loss from this region in KO tissues (Fig. 3E); brush border to cytosol ratios confirmed the redistribution of VAMP4 in the absence of PACSIN2 (1.86 ± 0.52 WT vs. 1.31 ± 0.45 KO; Fig. 3F). We also examined the localization of RAB14, another factor implicated in endocytic trafficking at the apical membrane of polarized epithelial cells [51]. Once again, this marker demonstrated decreased apical localization in the PACSIN2 KO tissue and a decreased brush border to cytosol ratio (Fig. S2). Thus, in addition to disrupting F-actin assembly throughout the enterocyte, these results show that loss of PACSIN2 disrupts the normal enrichment of endocytic machinery including Dynamin2, VAMP4, and RAB14, in the sub-apical terminal web.

**Figure 3.**
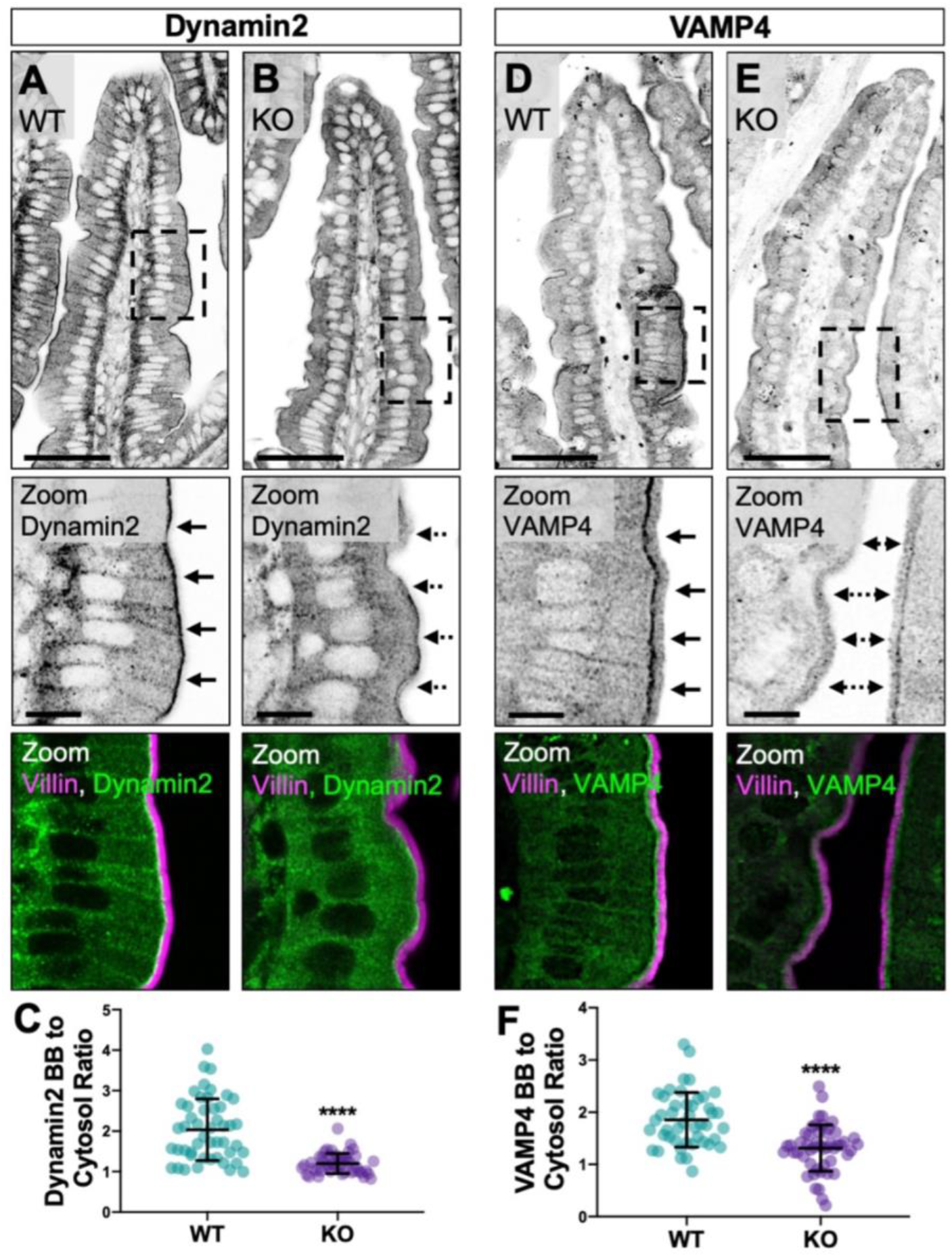
Endocytic machinery is mislocalized in the absence of PACSIN2. **(A, B)** Single confocal image planes of WT and PACSIN2 KO paraffin-embedded tissue sections stained with anti-Villin (magenta) to highlight the brush border and anti-Dynamin2 (green). Solid arrows in zoom panels highlight Dynamin2 signal at the base of the brush border in WT tissue (A), dashed arrows highlight mislocalization of Dynamin2 signal in KO tissue (B); Scale bars, 50 μm for main panel, 10 μm for zoom. (C) Quantification of the ratio of Dynamin2 brush border (BB) to cytosol signal intensity between WT and PACSIN2 KO (n = 48 measurements). (D, E) Single confocal image planes of WT and PACSIN2 KO paraffin-embedded tissue sections stained with anti-Villin (magenta) and anti-VAMP4 (green). Solid arrows in zoom panels highlight VAMP4 signal at the base of the brush border in WT tissue (D), dashed arrows highlight mislocalization of VAMP4 signal in KO tissue (E); Scale bars, 50 μm for main panel, 10 μm for zoom. (F) Quantification of the ratio of VAMP4 brush border (BB) to cytosol signal intensity between WT (n = 45 measurements) and PACSIN2 KO (n = 50 measurements). Error bars indicate ± SD; p values were calculated using a t test (****p<0.0001).

### Loss of PACSIN2 disrupts microvillar ultrastructure and organization

To understand how loss of PACSIN2 impacts brush border architecture, we employed transmission electron microscopy (TEM) to visualize WT and PACSIN2 KO tissues at the ultrastructural level (Fig. 4A,B). TEM imaging of sections parallel to the microvillar axis allowed us to perform detailed morphometry. Strikingly, microvilli in PACSIN2 KO brush borders were significantly shorter relative to WT (1.93 ± 0.35 μm WT vs. 1.08 ± 0.31 μm KO; Fig. 4C). We also found that the extent of membrane coverage, calculated as the percent of core actin bundle enveloped in membrane, was significantly reduced in KO brush borders (80.22 ± 3.16% WT vs. 66.34 ± 7.10% KO; Fig. 4B,D). Reduced membrane coverage was also linked to longer rootlets (0.47 ± 0.12 μm WT vs. 0.54 ± 0.15 μm KO; Fig. 4E). In addition, we noted a much more irregular membrane profile in the intermicrovillar region (Fig. 4F,G). In KO enterocytes, the straightness of this profile was significantly reduced compared to WT controls (0.87 ± 0.08 μm WT vs. 0.72 ± 0.11 μm KO; Fig. 4H). Upon closer inspection of the PACSIN2 KO terminal web, we found an increased number of membrane invaginations, most likely stalled endocytic intermediates, extending into the cytoplasm (1.98 ± 1.07 WT vs. 4.25 ± 1.87 KO; Fig. 4I). Combined with our staining data, these results indicate that loss of PACSIN2 disrupts endocytosis, which is associated with profound effects on microvillar morphology and the extent of membrane coverage.

**Figure 4.**
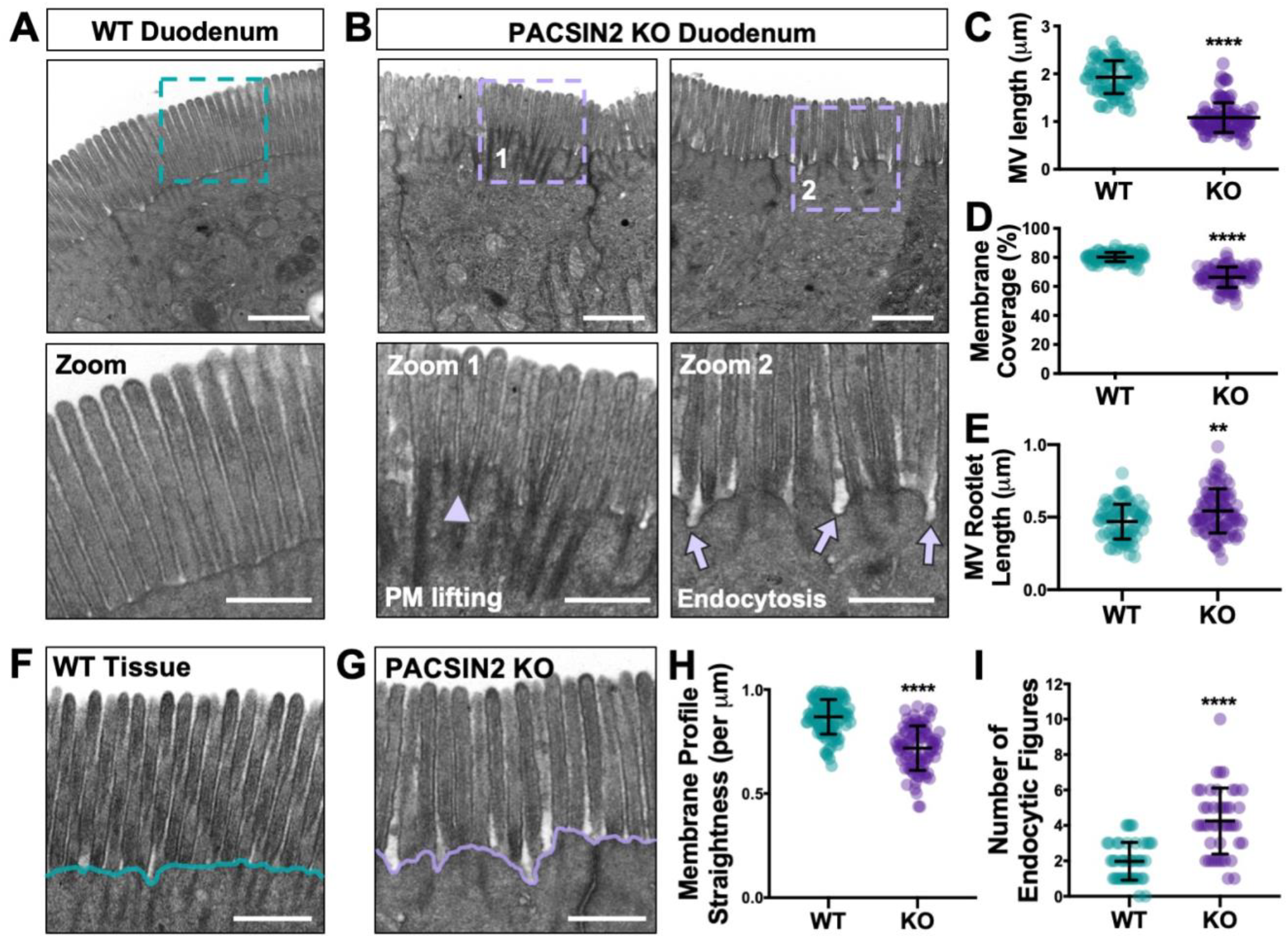
Loss of PACSIN2 disrupts microvillar ultrastructure and organization. **(A)** TEM image of a WT **brush border** in a plane parallel to the microvillar axis; teal dashed box indicates region highlighted in zoom panel below. Scale bar, 1 μm. **(B)** TEM images of PACSIN2 KO brush borders in a plane parallel to the microvillar axis; purple dashed boxes indicate region highlighted in zoom panels below. Arrowhead in zoom 1 highlights membrane lifting, arrows in zoom 2 highlight endocytic events. Scale bars, 1 μm. **(C)** Quantification of microvillar length in WT (n = 82) and KO (n = 102); measurements were selected so that only protrusions with actin cores fully visible along their length were scored. **(D)** Quantification of membrane coverage, the percentage of an actin core wrapped in membrane, in WT (n = 83) and KO (n = 102) microvilli. **(E)** Quantification of microvillar rootlet length in WT (n = 83) and KO (n = 102) microvilli. **(F, G)** Representative images of WT (F) and KO (G) tissue used in the quantification of membrane profile straightness (H); scale bars, 0.5 μm. Teal and purple lines highlight the decreased membrane straightness in KO. **(H)** Quantification of plasma membrane profile straightness at the base of WT (n = 102) and KO (n = 88) microvilli; total membrane length was measured over 1 μm units**. (I)** Quantification of the number of endocytic events, or structures that resemble stalled endocytic intermediates, at the plasma membrane of WT (n = 44 image fields) and KO (n = 44 image fields) brush borders. Error bars indicate ± SD; p values were calculated using a t test (**p<0.01, ****p<0.0001).

To further analyze the organization of PACSIN2 KO brush borders, we performed SEM to visualize the apical surface. *En face* images immediately revealed perturbations in microvillar packing, with more apparent free space between adjacent protrusions (Fig. S3). We also examined inter-microvillar spacing by calculating nearest neighbor distances (NND) for large numbers of protrusions. KO brush borders exhibited greater NND values with higher variability relative to WT controls (116.9 ± 14.6 nm WT vs. 131.8 ± 18.4 nm KO; Fig. S3C). To examine the impact of this increase in NND on the organization of microvilli, we calculated fast Fourier transforms (FFTs) as previously described [21]. FFTs generated by WT brush borders exhibited the expected hexagonal pattern with six prominent first order spots (Fig. S3D). However, FFTs generated from KO brush borders produced a pattern that lacked first order spots, indicating a loss of ordered packing (Fig. S3E). These data reveal that microvilli in the PACSIN2 KO brush borders are less densely packed and no longer organized in the hexagonal arrays that are a defining feature of normal enterocyte brush borders.

### Inhibition of endocytosis reduces microvillar membrane coverage

Our measurements indicate that under normal conditions, the distal ~80% of a microvillus actin core bundle is enveloped in apical plasma membrane (Fig. 4D). In the absence of PACSIN2, membrane coverage is significantly reduced with values that are much more variable across a population of protrusions (Fig. 4D). By promoting endocytic activity and/or anchoring the intermicrovillar membrane to the actin cytoskeleton, PACSIN2 could play a direct role in controlling the extent of microvillar membrane coverage. Because mechanisms that control microvillar membrane coverage remain poorly defined, we sought to test this hypothesis using the Ls174T-W4 (W4) intestinal epithelial cell culture model, which has been engineered to form microvilli upon exposure to doxycycline [52]. Similar to WT intestinal tissue, W4 cells demonstrate localization of PACSIN2 and Dynamin2 in the terminal web (Fig. S4A,B).

We first sought to determine if PACSIN2 KD in W4 cells generated a phenotype similar to what we observed with PACSIN2 KO mouse intestinal tissues. W4 cells transduced with scramble control shRNA or shRNA targeting PACSIN2 were fixed and stained to label the plasma membrane and underlying actin cytoskeleton, and then imaged using super-resolution structured illumination microscopy (SIM). KD of PACSIN2 significantly decreased microvillar membrane coverage relative to scramble controls (78.3 ± 8.6% KD vs. 89.1 ± 6.2% SCR) (Fig. 5A-C). We also imaged PACSIN2 KD W4 cells live using spinning disk confocal microscopy (SDCM). Remarkably, time-lapse acquisitions revealed the formation of long aberrant membrane tubules, presumably stalled endocytic intermediates, which originated in the intermicrovillar region (Fig. 5E). Coincident with the formation of these tubules, we noted significant apical membrane lifting, which exposed the rootlets of adjacent microvillar core actin bundles, in a manner that was strikingly reminiscent of membrane coverage perturbations observed in PACSIN2 KO brush borders (Fig. 4). Thus, in terms of the microvillar membrane coverage, PACSIN2 KD in W4 cells phenocopies the defects observed in brush borders from PACSIN2 KO mice.

**Figure 5.**
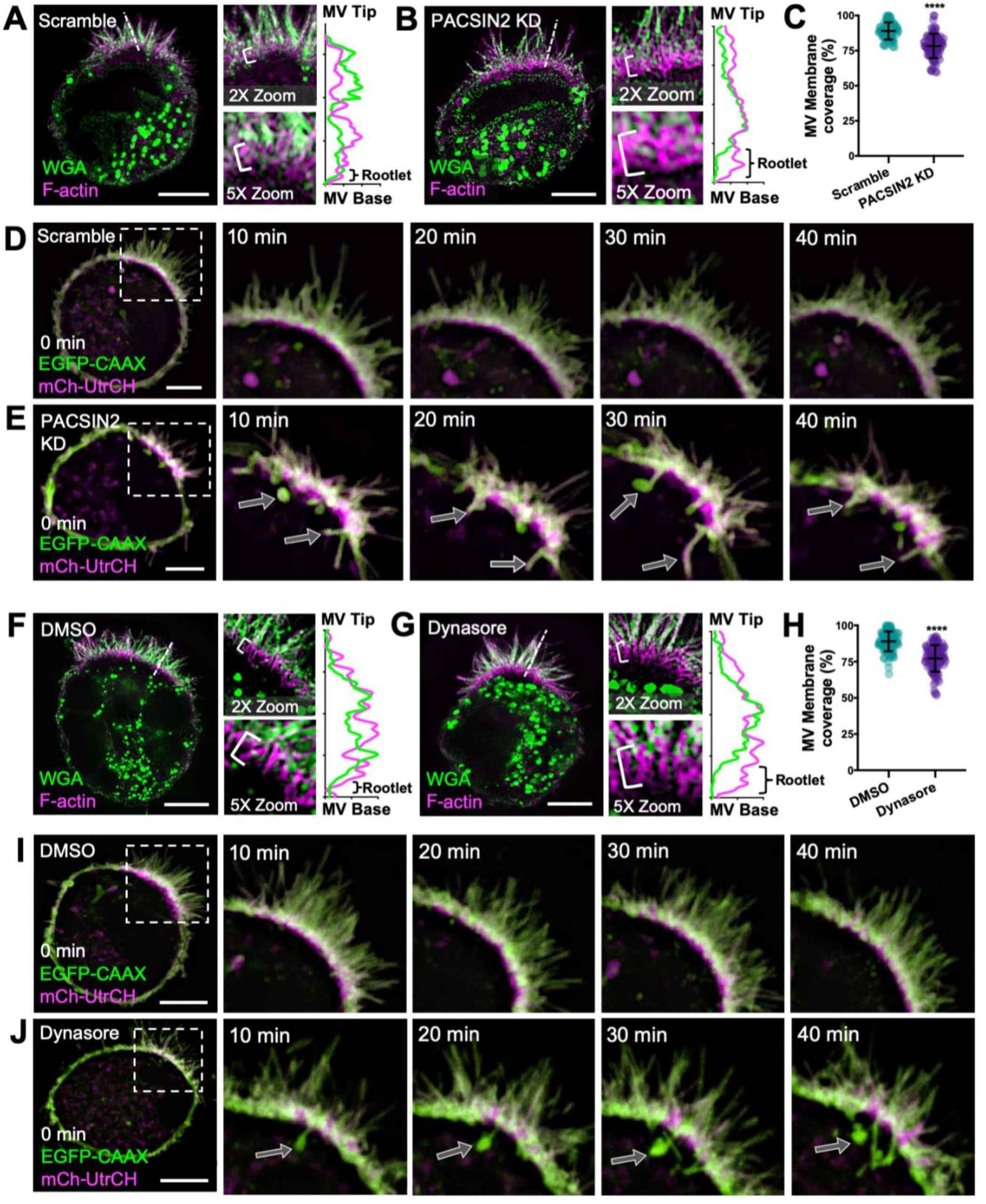
Inhibition of endocytosis reduces microvillar membrane coverage. **(A, B)** SIM projections of scramble control (A) and PACSIN2 KD (B) W4 cells stained for WGA (membrane, green) and phalloidin (magenta). Brackets in zoom panels indicate actin rootlet lengths. Dashed lines denote where line scans were drawn through a single microvillus to show increased actin rootlet length; membrane (green), actin (magenta). Scale bars, 5 μm. **(C)** Quantification of microvillar membrane coverage; scramble n = 77 microvilli from 10 cells; PACSIN2 KD n = 88 microvilli from 11 cells. **(D, E)** Montages of scramble control and PACSIN2 KD W4 cells expressing EGFP-CAAX box (last 10aa of the GTPase HRas; membrane, green) and mCherry-UtrCH (F-actin, magenta). Arrows in the PACSIN2 KD cell (E) indicate membrane tubules forming into the cytosol. Scale bars, 5 μm. **(F, G)** SIM projections of DMSO control (F) and 80 μM Dynasore (G) treated W4 cells stained for WGA (membrane, green) and phalloidin (magenta). Brackets in zoom panels indicate actin rootlet lengths. Dashed lines denote where line scans were drawn to show increased actin rootlet length; membrane (green), actin (magenta). Scale bars, 5 μm. **(H)** Quantification of microvillar membrane coverage; DMSO n = 104 microvilli from 13 cells; Dynasore n = 105 microvilli from 12 cells. **(I, J)** Montages of DMSO control and 80 μM Dynasore treated W4 cells expressing EGFP-CAAX box (membrane, green) and mCherry-UtrCH (F-actin, magenta). Arrows in the Dynasore treated cell (J) indicate membrane tubules forming into the cytosol. Scale bars, 5 μm. Error bars indicate ± SD; p values were calculated using a t test (****p<0.0001).

We next set out to determine if the microvillar membrane coverage defects observed in PACSIN2 KO tissues and KD W4 cells were due specifically to perturbations in endocytic activity. For these experiments, we exposed differentiating W4 cells to Dynasore, a small molecule inhibitor of the GTPase domain of Dynamin2 that is expected to prevent the scission of endocytic vesicles from the apical membrane. Dynasore-treated W4 cells were fixed and stained to visualize the actin cytoskeleton and plasma membrane, and then imaged using SIM. Remarkably, exposure to Dynasore decreased microvillar membrane coverage significantly relative to control DMSO-treated cells (77.2 ± 9.1% Dynasore vs. 89.0 ± 6.9% DMSO; Fig. 5F-H). We also used SDCM to image the impact of Dynasore treatment on live W4 cells. Similar to that observed in PACSIN2 KD W4 cells, we noted the formation of long aberrant membrane tubules, which again originated in the intermicrovillar region (Fig. 5J). The formation of these tubules also coincided with significant membrane lifting and exposure of microvillar core bundle rootlets (Fig. 5J). We verified this effect using a second inhibitor of endocytosis, Pitstop 2, which generated similar aberrant tubule formation and membrane lifting (Fig. S4D). Together, these findings uncover a previously unrecognized link between PACSIN2-dependent endocytic activity and the extent of microvillar membrane coverage. These data further suggest that inward forces on the apical membrane, normally generated by endocytic machinery, serve to control microvillar morphology.

## DISCUSSION

PACSIN family proteins have long been implicated in the regulation of actin assembly in the context of membrane deformation during endocytosis and vesicle formation. Indeed, in the initial report, PACSIN1 (primarily expressed in neural tissues) co-immunoprecipitated with synaptic vesicle endocytic factors including dynamin, synaptojanin, and synapsin-1, as well as N-WASP, an actin nucleation promoting factor that activates the ARP2/3 complex [53]. All of these interactions were mediated through the PACSIN1 C-terminal SH3 domain [53]. During endocytosis, PACSINs are believed to recruit N-WASP, which in turn targets ARP2/3 to generate bursts of actin filament polymerization in the space between the plasma membrane and nascent budding vesicles. Combined with activity of the Dynamin GTPase, which constricts the necks of forming vesicles, these bursts of actin polymerization likely generate additional mechanical force for efficient vesicle scission [43, 48]. Although PACSINs have been implicated in various forms of endocytosis, including activity dependent bulk endocytosis (ADBE) and clathrin-mediated endocytosis [36], PACSIN2 has more recently been implicated in caveolar endocytosis, where it binds to the necks of nascent caveolae and recruits Dynamin2 to promote vesicle scission [33, 34, 46]. In support of an endocytic role in transporting epithelia, previous studies localized PACSIN2 to the sub-apical terminal web region of native enterocytes in the mouse small intestine and human W4 cells in culture [6]. In the terminal web, endocytic vesicles are formed from the inwardly curving membrane found between neighboring microvilli [54]. Indeed, SIM imaging of differentiated W4 cells revealed robust PACSIN2 localization in the intermicrovillar region, immediately between adjacent core actin bundles [6]. In the present study, we found that markers of endocytosis which are normally enriched in the terminal web, including Dynamin2, VAMP4, and RAB14, were also lost from this region in the absence of PACSIN2. Together, all of these data establish a role for PACSIN2 in the normal targeting of endocytic machinery to the sub-apical compartment.

In addition to a role in apical endocytic vesicle formation, PACSIN2 was also found to play a role in recruiting the linear actin nucleator, COBL, to the terminal web. In the W4 cell culture model, COBL loss-of-function impairs brush border assembly, whereas overexpression promotes the formation of microvillar actin cores in a manner that depends on the number WH2 domains [6, 7]. COBL is also recruited to the apex of epithelial cells coincident with the earliest events in brush border assembly [6]. Consistent with its role in targeting COBL to the terminal web, we found significantly lower levels of COBL at the base of the brush border in PACSIN2 KO tissues. PACSIN2 KO enterocytes also exhibited reduced apical actin levels as assessed with F-actin reporter, phalloidin (Fig. 2). Confocal volume projections showed a clear thinning of brush border F-actin signal, with reduced microvillar density and regions that appeared to lack microvilli completely (Fig. 2C,E). In the ultrastructural analysis of KO tissues, we also noted a significant decrease in microvillar length (Fig. 4C). Together these findings suggest that KO of PACSIN2, and subsequent loss of COBL from the terminal web, impairs the production of actin filaments that form microvillar actin core bundles.

Remarkably, measurements of phalloidin intensity from other parts of PACSIN2 KO enterocytes revealed lower levels of F-actin, although the relative ratio of apical/cell body F-actin signal remained unchanged in response to PACSIN2 KO (Fig. 2K). Because most of the cell body signal derives from the basolateral margins, we propose that these perturbations are induced by loss of N-WASP stimulated ARP2/3 activity at the basolateral cortex. In support of this, previous studies showed that inactivation of ARP2, a component of the ARP2/3 complex, decreased actin polymerization and impairs the morphology and stability of epithelial adherens junctions [55, 56]. Inhibition of actin polymerization also impairs adherens junction reassembly and reduces E-cadherin enrichment [57–59]. Interestingly, in our studies, the loss of junctional actin correlates with the loss of ZO-1 and E-Cadherin signal in the PACSIN2 KO mouse (Fig. S4), indicating a disruption in normal junctional stability and architecture.

Perhaps the most unexpected finding from the current investigation was the striking perturbation in microvillar ultrastructure in PACSIN2 KO brush borders. In PACSIN2 KO brush borders, we observed a significant decrease in microvillar length and the extent of membrane coverage, i.e. the fraction of core actin bundle encapsulated in plasma membrane. These changes were also accompanied by a corresponding increase in the length of exposed rootlet. How does loss of PACSIN2 impact microvillar structure and membrane coverage? While it is known that membrane-cytoskeleton linkers, such as Myo1a and Ezrin, stabilize physical contact between the plasma membrane and the underlying actin core, factors that control the extent of membrane coverage are poorly understood. A clue to the mechanism might come from our observation of a higher frequency of membrane invaginations originating from the intermicrovillar region in PACSIN2 KO brush borders. Because PACSIN2 and its binding partners (e.g. Dynamin) normally stimulate vesicle scission at these sites, the elongated invaginations that extend through the terminal web are most likely stalled endocytic structures, an interpretation consistent with their tubular morphology. Indeed, PACSIN2 KD in cultured cells has been shown to generate elongated caveolae [33]. If the entire apical membrane is composed of a single continuous surface, the formation of exaggerated tubules in the terminal web would directly reduce the amount of membrane material available for encapsulating microvilli and thus, compromise the extent of membrane coverage. To test this possibility more directly, we modeled the defects observed in PACSIN2 KO tissues in the W4 intestinal epithelial cell line. PACSIN2 KD in this context also lead to reduced membrane coverage of microvilli. Strikingly, we also observed that the inward pulling of exaggerated tubules temporally coincides with loss of membrane coverage on microvilli immediately adjacent to these sites. Because we were able to phenocopy these events with two distinct inhibitors of endocytosis, we conclude that the exaggerated tubules observed in PACSIN2 KO and PACSIN2 KD cells are in fact stalled endocytic intermediates. Together our findings highlight a mechanistic link between sub-apical endocytic activity and the membrane coverage of apical microvilli.

Interestingly, a role for inward pulling forces on the apical plasma membrane in shaping finger-like protrusions has been highlighted in previous studies of the pointed-end directed motor, MYO6. MYO6 localizes to the terminal web where it interacts with endocytic machinery near the pointed-ends of microvillar core actin bundles, including DAB2, and GIPC [60]. In *Snell’s Waltzer* mice, which lack functional MYO6, inner ear hair cells exhibit a membrane lifting phenotype similar what we observe in PACSIN2 KO brush borders [61]. These cells also manifest with fused or coalesced protrusions, where multiple core bundles appear to be enveloped in a single tubule of plasma membrane. Later studies with the same model system revealed similar phenomena in the enterocyte brush border, with marked decreases in the membrane coverage of core actin bundles and more general disorder in the terminal web [62]. In combination with the data we present here, these studies lead to a model whereby the formation and steady-state morphology of finger-like protrusions such as microvilli and stereocilia, are controlled by a balance of outward and inward mechanical forces that impinge on the plasma membrane. PACSIN2 likely limits these forces by promoting the budding and scission of endocytic vesicles from the intermicrovillar membrane. Whether PACSIN2 functions in the same pathway as MYO6 is not known, but functional links between these two factors should be the focus of future studies.

## MATERIALS AND METHODS

### Frozen Tissue Preparation

Segments of WT and KO intestine were removed and flushed with PBS and pre-fixed for 10 minutes with 4% paraformaldehyde (PFA) to preserve the tissue structure. The tube was then cut along its length, sub dissected into 0.5 μm square chunks, fixed for an additional 30min in 4% PFA at RT, and washed 3 times in PBS. Samples were then gently placed on top of a 30% sucrose solution in TBS and allowed to sink to the bottom overnight at 4°C. Specimens were then swirled in three separate blocks of OCT (Electron Microscopy Sciences), oriented in a block filled with fresh OCT, and snap-frozen in dry ice-cooled acetone. Samples were cut in 10 μm sections and mounted on slides for staining.

### Cell Culture

Ls174T-W4 cells (female *Hs* colon epithelial cells) were cultured in DMEM with high glucose and 2 mM L-glutamine supplemented with 10% tetracycline-free fetal bovine serum (FBS), G418 (1 mg/ml), blasticidin (10 μg/ml), and phleomycin (20 μg/ml). The cell line was obtained from Dr. Hans Clevers (Utrecht University, Netherlands) and has not been additionally authenticated. All cells were grown at 37°C and 5% CO2.

### Transfections and lentivirus production

All transfections were performed using Lipofectamine 2000 (Invitrogen) according to the manufacturer’s instructions and the cells were allowed to recover overnight (ON). Lentivirus was generated by co-transfecting HEK293FT cells (Fetal *Hs* embryonic epithelial cells; T75 flasks at 80% confluency) with 6 μg of pLKO.1 PACSIN2 shRNA KD plasmids (Open Biosystems; TRCN0000037980), 4 μg of psPAX2 packaging plasmid, and 0.2 μg of pMD2.G envelope plasmid using FuGENE 6 (Promega). Cells were incubated up to 48 hrs and then lentivirus-containing media was collected and concentrated with Lenti-X concentrator (Clontech). To transduce W4 cells in T25 flasks, lentiviral shRNAs with 6 μg/ml polybrene (Sigma) was added dropwise to the media. After a 24-hour incubation, the media was changed and resupplemented with lentiviral shRNAs for an additional 24 hours. The cells were then seeded into 6-well plates with glass coverslips and incubated ON in the absence or presence of 1 μg/ml doxycycline, and then prepared for immunofluorescence.

### Immunofluorescence

Frozen tissue sections of WT and PACSIN2 KO intestinal tissue were washed in phosphate-buffered saline (PBS) three times and permeabilized for 10 min with 0.1% Triton X-100/PBS at RT. The tissue sections were then blocked with 10% bovine serum albumin (BSA) at 37°C for 2 hours and washed once with PBS. Primary antibodies (listed below) were diluted in 10% BSA/PBS and incubated with cells at 4°C O/N, followed by four washes with PBS. Tissue sections were then stained with phalloidin and secondary antibodies (listed below) in 1% BSA/PBS for 2 hrs at RT, washed three times with PBS and mounted with Prolong Gold Antifade mounting media (P36930; Invitrogen). Paraffin-embedded small intestinal tissue sections of WT and PACSIN2 KO were deparaffinized using Histo-clear solution (Fisher) and rehydrated in a descending graded ethanol series. Slides were then subject to an antigen retrieval step consisting of boiling for 1 hr in a solution of 10 mM Tris (pH 9.0) and 0.5 mM EGTA. Slides were then washed in PBS three times and stained O/N at 4°C with primary antibodies (see below) in 10% BSA/PBS. After washing with PBS four times, samples were stained with secondary antibodies in 1% BSA/PBS for 2 hrs at RT. Slides were then washed four times with PBS and mounted in ProLong Gold Antifade mounting media.

For SIM imaging, cells were plated on glass coverslips and allowed to adhere for at least 6 hrs. They were then washed with pre-warmed PBS and fixed with warm 4% PFA/PBS for 15 min at 37°C. Cells were then washed three times with PBS and permeabilized with 0.1% Triton X-100/PBS for 15 min at RT. Cells were once again washed three times with PBS and blocked for 1 hr at 37°C in 5% BSA/PBS. Primary antibodies (listed below) were diluted in 1% BSA/PBS and incubated with cells at 37°C for 1 hr, followed by four washes with PBS. Cells were then incubated for 1 hr with secondary antibodies and phalloidin (listed below) at RT. Coverslips were then washed four times with PBS and mounted on glass slides in ProLong Gold Antifade Mounting Media. For live cell spinning disk confocal imaging of W4 cells, previously transfected cells were plated on glass-bottom dishes with 1 μg/ml of doxycycline and allowed to adhere for 6 hours. If drug treatments were performed, 80μM DMSO/ 80μM Dynasore (D7693; Sigma-Aldrich) or 30μM DMSO/ 30μM Pitstop 2 (SML1169; Sigma-Aldrich) were diluted into 1ml media and added to glass-bottom dish of W4s 10 minutes before acquisition. For live imaging of scramble/ PACSIN2 KD cells, the protocol above was used however cells were seeded into glass-bottom dishes instead of 6-well plates and induced with 1μg/ml doxycycline and allowed to adhere for at least 4-6 hours before imaging. Movies of single W4 cells were acquired every 5 seconds for 30 minutes or continuously for 4 minutes. All live cells were maintained in a humid environment at 37°C and 5% CO2 using a stage-top incubation system. Image acquisition was controlled with Nikon Elements software.

The following dilutions were used for primary antibodies for staining: anti-PACSIN2 (2.5 μg/ml, HPA049854; Sigma-Aldrich), anti-COBL (1μg/ml, HPA019033; Sigma-Aldrich), anti**-** Dynamin2 (4 μg/ml, NBP2-47477; Novus Biologicals), anti-villin (4μg/ml; Santa Cruz #sc-66022), anti-E-Cadherin (0.5 μg/ml; BD Biosciences #610182), anti-ZO-1 (5μg/ml, 61-7300; Thermo Fisher), anti-VAMP4 (2μg/ml, HPA050418; Sigma-Aldrich), anti-RAB14 (4μg/ml; Invitrogen #PA5-55306). The following dilutions were used for secondary antibodies and cell dyes for staining: goat anti-rabbit Alexa Fluor 488 F(ab’)2 Fragment (2 μg/ml, A11070; Molecular Probes), goat anti-mouse Alexa Fluor 488 F(ab’)2 Fragment (2 μg/ml, A11017; Molecular Probes), Alexa Fluor 568–phalloidin (1:200, A12380; Invitrogen), or Wheat Germ Agglutin Oregon Green (WGA) (2μg/ml, W67-48; Life Technologies).

### Light Microscopy

Confocal microscopy was performed using a Nikon A1R laser-scanning confocal microscope equipped with 60x/1.4 NA and 100x/1.49 NA objectives. SIM was performed using a Nikon N-SIM with an Apo TIRF 100x/1.49 NA objective. All images used for quantitative comparisons were prepared with equal treatment, acquired with identical parameters (e.g. pinhole diameter, detector gain), and processed in an identical manner. Richardson-Lucy deconvolution of image volumes (20 iterations) was performed using Nikon Elements software. Live-cell imaging of W4 cells was performed on a Nikon Yokogawa CSU-X1 spinning disk confocal microscope. Images were contrast enhanced and cropped using ImageJ software (NIH).

### Electron Microscopy

Segments of WT and KO intestine were placed into 0.1M HEPES (pH 7.3) and sub dissected into 2mm chunks at RT. Samples were placed into scintillation vials and incubated in RT fix buffer (4% PFA, 2.5% glutaraldehyde, 2mM CaCl_2_ in 0.1M HEPES) for 1 hr and washed 3 times in HEPES buffer. Samples were incubated with 1% tannic acid/HEPES for 1 hr, washed 3 times with ddH_2_O followed by incubation with 1% osmium tetroxide/ddH_2_O for 1 hour. Samples were then washed 3 times with ddH_2_O, incubated in 1% uranyl acetate/ddH_2_O for 30 min then washed with ddH_2_O. Samples were dehydrated in a graded ethanol series and then dried using critical point drying. Samples were then mounted on aluminum stubs and coated with gold/palladium using a sputter coater. Imaging was performed using a Quanta 250 Environmental SEM operated in high vacuum mode with an accelerating voltage of 5 kV. All EM reagents were purchased from Electron Microscopy Sciences.

### Image Analysis and Statistics

All image analysis and signal intensities measurements from image data were performed using FIJI or Nikon Elements software. To perform intensity analyses (Figures 1, 2 and 6), the brush border and/ or cytosol were thresholded in villar confocal images using Nikon Elements software and the mean intensity numbers per villus were plotted; brush border to cytosol enrichment was defined as the ratio of these two mean intensities. Microvillar length measurements were performed on projected SIM images (Supplemental Figure 3) or on TEM images (Figure 4) by tracing individual microvillar actin bundles using FIJI. For W4 cell microvillar length analysis, at least 10 microvillar actin bundles were scored per cell and at least 25 cells measured per experiment. Microvillar membrane coverage measurements were performed on projected W4 SIM images (Figure 5) or on TEM images by dividing the length of a microvilli covered in membrane by the entire actin bundle from the rootlet to the tip. Nearest neighbor distance measurements (Figure 3) were performed by thresholding microvilli in SEM images using Nikon Elements. Data were analyzed with a D’Agostino and Pearson omnibus normality test to determine normal distribution and normally distributed data were statistically analyzed to determine significance using the unpaired Student’s *t* test. Welch’s correction was used in cases where data sets did not exhibit equal variance. Statistical analyses performed are stated in the figure legends. All graphs were generated and statistical analyses performed using Prism (v.7, GraphPad).

### Animal Studies

Animal experiments were carried out in accordance with Vanderbilt University Medical Center Institutional Animal Care and Use Committee guidelines.

## Supporting information

Postema et al Supp Figures

## MOVIE LEGENDS

**Movie S1, Related to Figure 5. Live imaging of Ls174T-W4 cell treated with 80 μM DMSO.**

Spinning disk confocal imaging of an induced Ls174T-W4 cell expressing EGFP-CAAX (green, membrane) and mCherry-UtrCH (magenta, F-actin). Cell was treated with 80 μM DMSO ~10 min prior to acquisition. Movie was acquired every 30 seconds for 60 minutes and is played at 12.5 FPS. Scale bar, 5 μm.

**Movie S2, Related to Figure 5. Live imaging of Ls174T-W4 cell treated with 80 μM Dynasore**.

Spinning disk confocal imaging of an induced Ls174T-W4 cell expressing EGFP-CAAX (green, membrane) and mCherry-UtrCH (magenta, F-actin). Cell was treated with 80 μM Dynasore ~10 min prior to acquisition. Movie was acquired every 30 seconds for 90 minutes and is played at 12.5 FPS. Scale bar, 5 μm.

**Movie S3, Related to Figure 5. Live imaging of scramble control shRNA Ls174T-W4 cell.**

Spinning Disk confocal imaging of an induced, scramble control Ls174T-W4 cell expressing EGFP-CAAX (green, membrane) and mCherry-UtrCH (magenta, F-actin). Movie was acquired every 30 seconds for 60 minutes and is played at 12.5 FPS. Scale bar, 5 μm.

**Movie S4, Related to Figure 5. Live imaging of PACSIN2 shRNA Ls174T-W4 cell.**

Spinning Disk confocal imaging of an induced, IRTKS KD Ls174T-W4 cell expressing EGFP-CAAX (green, membrane) and mCherry-UtrCH (magenta, F-actin). Movie was acquired every 30 seconds for 90 minutes and is played at 12.5 FPS. Scale bar, 5 μm.

**Movie S5, Related to Figure S4. Live imaging of Ls174T-W4 cell treated with 30 μM Pitstop 2.** Spinning disk confocal imaging of an induced Ls174T-W4 cell expressing EGFP-CAAX (green, membrane) and mCherry-UtrCH (magenta, F-actin). Cell was treated with 30 μM Pitstop 2 ~10 min prior to acquisition Movie was acquired every 30 seconds for 90 minutes and is played at 12.5 FPS. Scale bar, 5 μm.

