## Supplementary material for "PACSIN2-dependent apical endocytosis regulates the morphology of epithelial microvilli": Postema et al Supp Figures

**CONTAINS:**

**Supplemental Figures 1-4**
**Supplemental Figure Legends**

#### Figure S1 (Related to Figures 1 and 2)

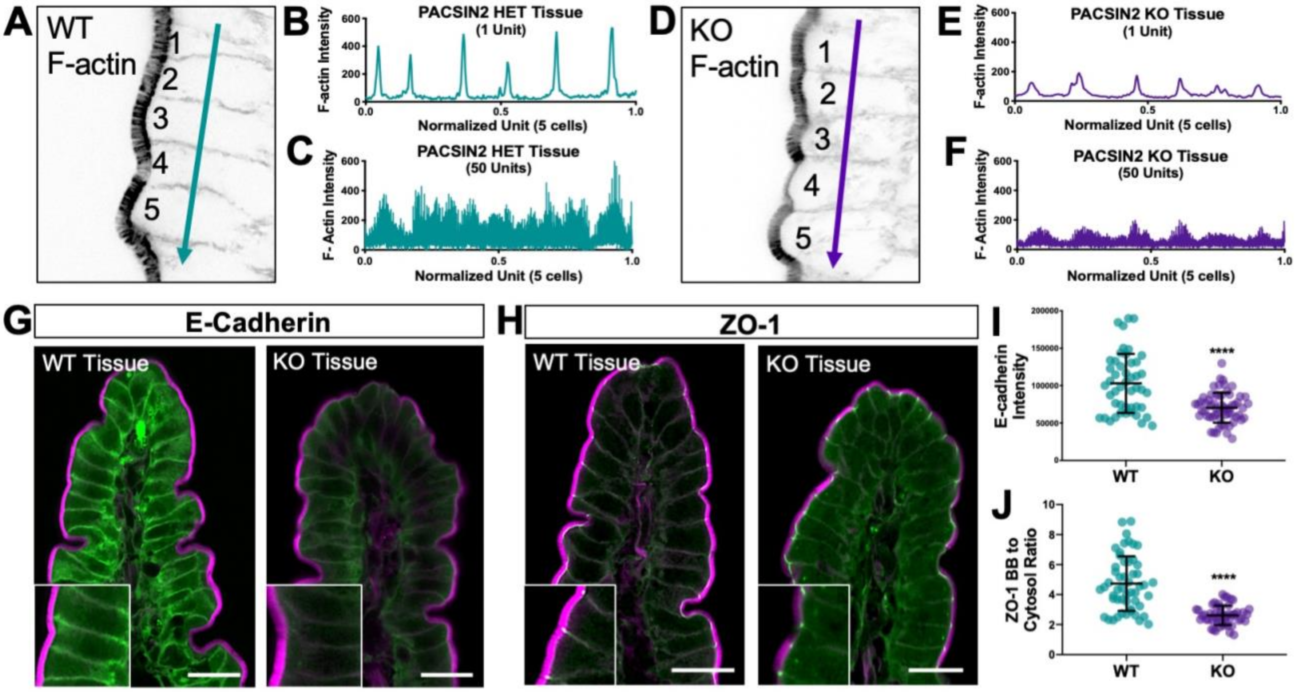


**Supplemental Figure 1, Related to Figure 2. Loss of PACSIN2 leads to junctional instability. (A, D)** Representative images used in quantification in D, E, G, H; phalloidin stained. **(B, C)** Top, raw intensity data of a line (depicted by teal arrow in C) through 5 total cells in WT tissue, peaks indicate the actin junctional intensity. Bottom, raw intensity data of lines through 5 cells in 50 WT tissue sections, lines have been smoothed for ease of viewing. **(E, F)** Top, raw intensity data of a line (depicted by purple arrow in F) through 5 cells in PACSIN2 KO tissue, peaks indicate the actin junctional intensity. Bottom, raw intensity data of lines through 5 total cells in 50 KO tissue sections, lines have been smoothed for ease of viewing. **(G)** Endogenous E-Cadherin (green) and phalloidin (F-actin, magenta) labelling of WT and PACSIN2 KO frozen tissue sections. Scale bars, 20 μm. **(H)** Endogenous ZO-1 (green) and phalloidin (F-actin, magenta) labelling of WT and PACSIN2 KO frozen tissue sections. Scale bars, 20 μm. **(I)** Quantification of E-Cadherin signal intensity between WT (n = 47 measurements) and PACSIN2 KO (n = 60 measurements). **(J)** Quantification of the ratio of ZO-1 BB to cytosol ratio between WT (n = 52 measurements) and PACSIN2 KO (n = 50 measurements). Error bars indicate ± SD; p value was calculated using a t test (****$p<0.0001$).

Figure S2 (Related to Figure 3)

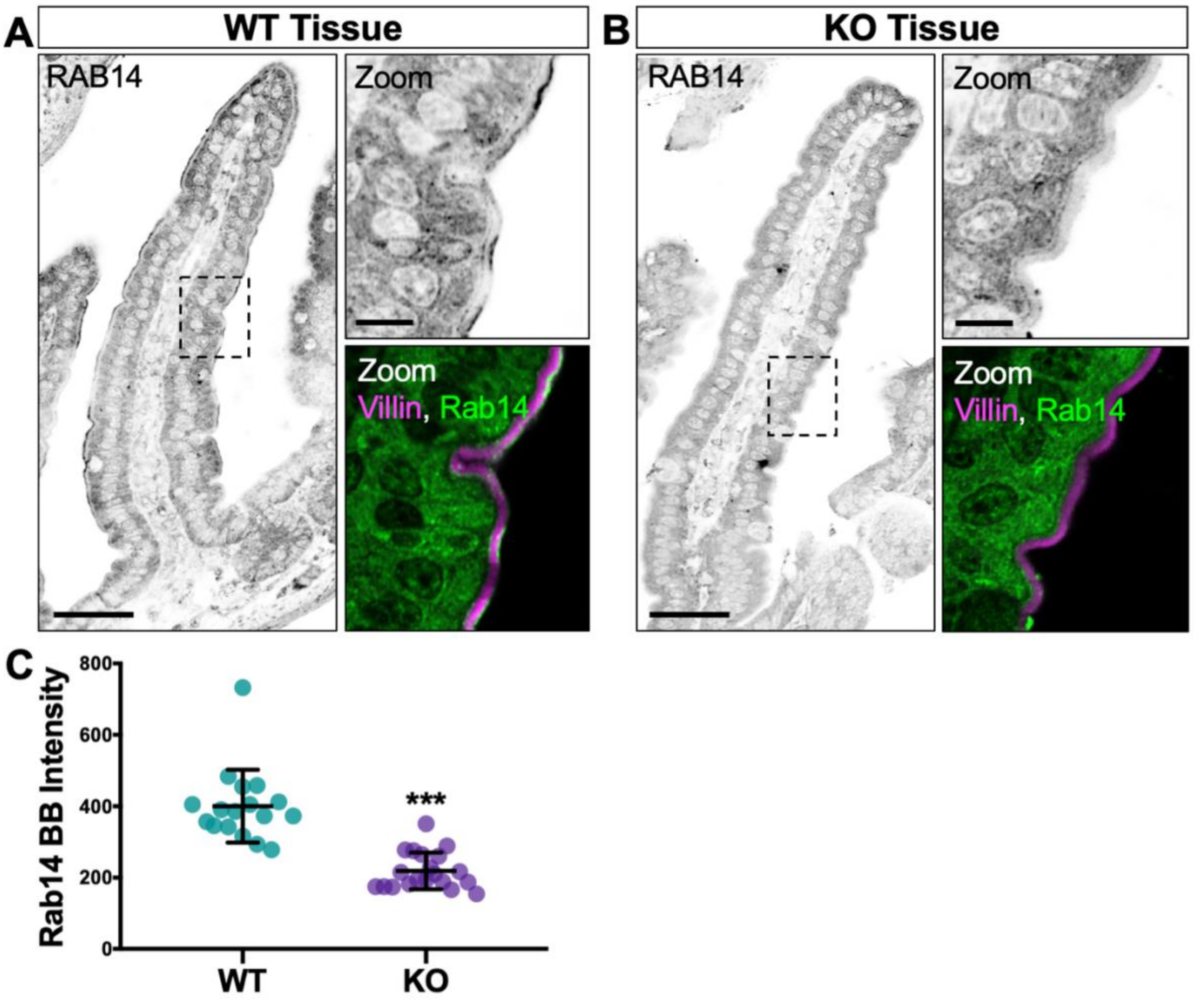


**Supplemental Figure 2, Related to Figure 3. Endocytosis marker Rab14 is mislocalized in the KO mouse. (A, B)** Single confocal image planes of WT and PACSIN2 KO paraffin-embedded tissue stained with anti-Villin (magenta) to highlight the brush border and anti-Rab14 (green). Signal is inverted for ease of viewing; scale bars, 50 μm in main panels, 10 μm in zoom. **(C)** Quantification of Rab14 signal intensity between WT (n = 17 measurements) and PACSIN2 KO (n = 20 measurements). Error bars indicate ± SD; p value was calculated using a t test (***$p<0.001$).

### Figure S3 – related to Figure 4

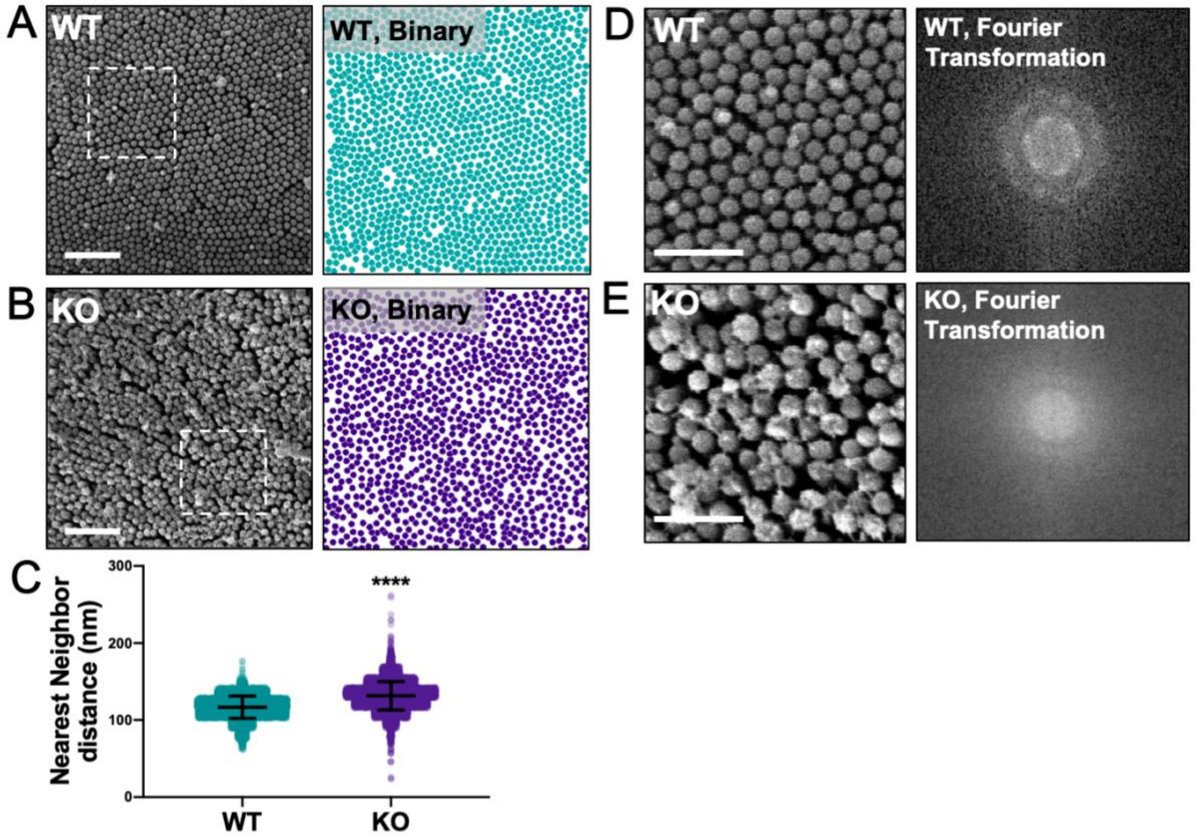


**Supplemental Figure 3, Related to Figure 4. Microvillar packing is decreased in the PACSIN2 KO mouse. (A, B)** SEM images WT (A) and KO (B) brush borders reveal microvillar packing defects in KO samples. Binary images indicating microvilli were used to determine the nearest neighbor distances. **(C)** Quantification of the nearest neighbor distance from mask. The mean center to center distance between microvilli was calculated; 6 fields of microvilli per condition. Error bars indicate ± SD; p value was calculated using a t test (****$p<0.0001$). **(D, E)** 1.5 $\mu m^2$ images showing differences in microvillar packing and the abnormal Fourier transformation obtained in the KO sample.

Figure S4 (Related to Figure 5)

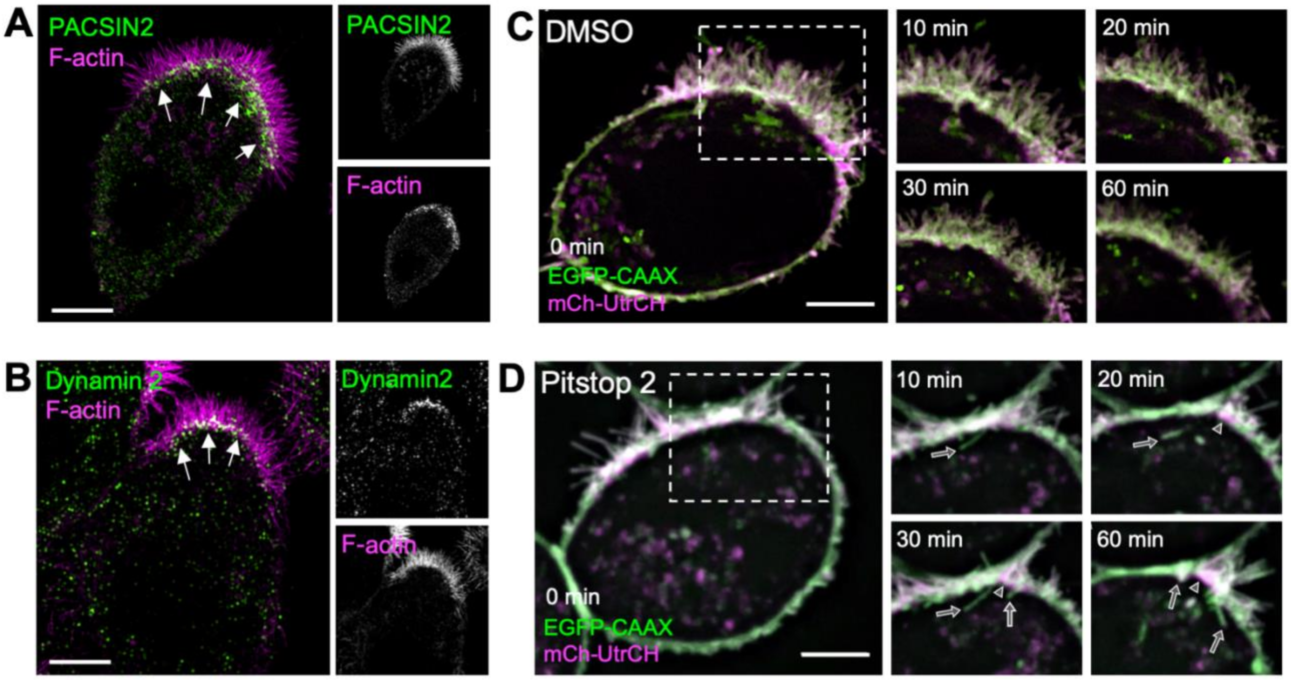


**Supplemental Figure 4, Related to Figure 5. Pitstop2 inhibits endocytosis in W4 cells. (A)** SIM projection of a W4 cell showing endogenous PACSIN2 (green) and stained with phalloidin (magenta). Arrows point to PACSIN2 puncta at the base of the brush border. **(B)** SIM projection of a W4 cell showing endogenous Dynamin2 (green) and stained with phalloidin (magenta). Arrows point to Dynamin2 puncta at the base of the brush border. **(C, D)** Montages of DMSO control and 30 μM Pitstop 2 treated W4 cell expressing EGFP-CAAX box (membrane, green) and mCherry-UtrCH (F-actin, magenta). Arrows in Pitstop 2 cell (D) indicate membrane tubules forming into the cytosol, arrowheads indicate membrane lifting. Scale bars, 5 μm.
